# NanoJ-SQUIRREL: quantitative mapping and minimisation of super-resolution optical imaging artefacts

**DOI:** 10.1101/158279

**Authors:** Siân Culley, David Albrecht, Caron Jacobs, Pedro Matos Pereira, Christophe Leterrier, Jason Mercer, Ricardo Henriques

## Abstract

Most super-resolution microscopy methods depend on steps that contribute to the formation of image artefacts. Here we present NanoJ-SQUIRREL, an ImageJ-based analytical approach providing a quantitative assessment of super-resolution image quality. By comparing diffraction-limited images and super-resolution equivalents of the same focal volume, this approach generates a quantitative map of super-resolution defects, as well as methods for their correction. To illustrate its broad applicability to super-resolution approaches we apply our method to Localization Microscopy, STED and SIM images of a variety of in-cell structures including microtubules, poxviruses, neuronal actin rings and clathrin coated pits. We particularly focus on single-molecule localisation microscopy, where super-resolution reconstructions often feature imperfections not present in the original data. By showing the quantitative evolution of data quality over these varied sample preparation, acquisition and super-resolution methods we display the potential of NanoJ-SQUIRREL to guide optimization of superresolution imaging parameters.

## Introduction

Super-resolution microscopy is a collection of imaging approaches achieving spatial resolutions beyond the diffraction limit of conventional optical microscopy (∼250 nm). Notably, methods based on Localization Microscopy (LM) such as Photo-Activated Localization Microscopy (PALM) (1) and Stochastic Optical Reconstruction Microscopy (STORM) (2) can achieve resolutions better than 30 nm. Due to their easy implementation and the large set of widely accessible resources developed by the research community these methods have become widespread (3–6). The quality and resolution achieved by super-resolution is largely dependent on factors such as the photophysics of fluorophores used (7), chemical environment of the sample (7, 8), imaging conditions (4, 5, 8, 9) and analytical approaches used to produce the final super-resolution images (9, 10). Balancing these factors is critical to ensure that the recovered data accurately represents the underlying biological structure. Thus far data quality assessment in super-resolution relies on researcher-based comparison of the data relative to prior knowledge of the expected structures (11, 12) or benchmarking of the data against other high-resolution imaging methods such as Electron Microscopy (1). An exception exists in the Structured Illumination Microscopy (SIM) field (13), where analytical frameworks exist for quantitative evaluation of image quality (14, 15). The simplest, most robust way to visually identify defects in super-resolution images is the direct comparison of diffraction-limited and super-resolved images of a sample. Assuming that the images represent the same focal volume, the super-resolution image should provide an improved resolution representation of the reference diffraction-limited image. While this allows for identification of common large scale artefacts, such as misrepresentation or disappearance of structures (16), details significantly smaller than the diffraction limit of the microscope cannot be validated. In addition as this analysis is performed empirically it is subject to human bias and interpretation.

Here we present SQUIRREL, a new analytical approach to quantitatively map local image errors and hence assist in their reduction. This is implemented as an easy-to-use, open-source ImageJ and Fiji (18) plugin (dubbed NanoJ-SQUIRREL), exploiting high-performance GPU-enabled computing. SQUIRREL is founded solely on the premise that a super-resolution image should be a high-precision representation of the underlying nanoscale position and photon emission of the imaged fluorophores. Although based on the principle of comparing conventional and super-resolution images, in contrast to other approaches, it requires no *a priori* knowledge of the expected structural properties of the sample or photophysics of the labels used. Therefore, assuming the imaged field-of-view has a spatially invariant point-spread-function (PSF), application of a resolution rescaling transfer function to the super-resolution image should produce an image with a high degree of similarity to the original diffraction-limited one. Variance between these images beyond a noise floor can be used as a quantitative indicator of local macro-anomalies in the super-resolution representation (Fig. 1). We show that, when using a conventional diffraction limited image as a reference, SQUIRREL is able to identify artefacts on scales ∼ 150 nm within a ∼ 600 nm depth focal volume. Identification of artefacts on a smaller scale can be achieved by cross-validations of different superresolution methods. This is demonstrated in LM, Stimulated Emission Depletion (STED) microscopy (19) and SIM. As a result we demonstrate that mapping of local errors can be used to compare, rank and identify discrepancies between super-resolution images from different methods. By fusing multiple super-resolution images we further show that it is possible to minimize reconstruction algorithm-specific artefacts, while providing insight into optimal super-resolution imaging conditions.

**Fig. 1.**
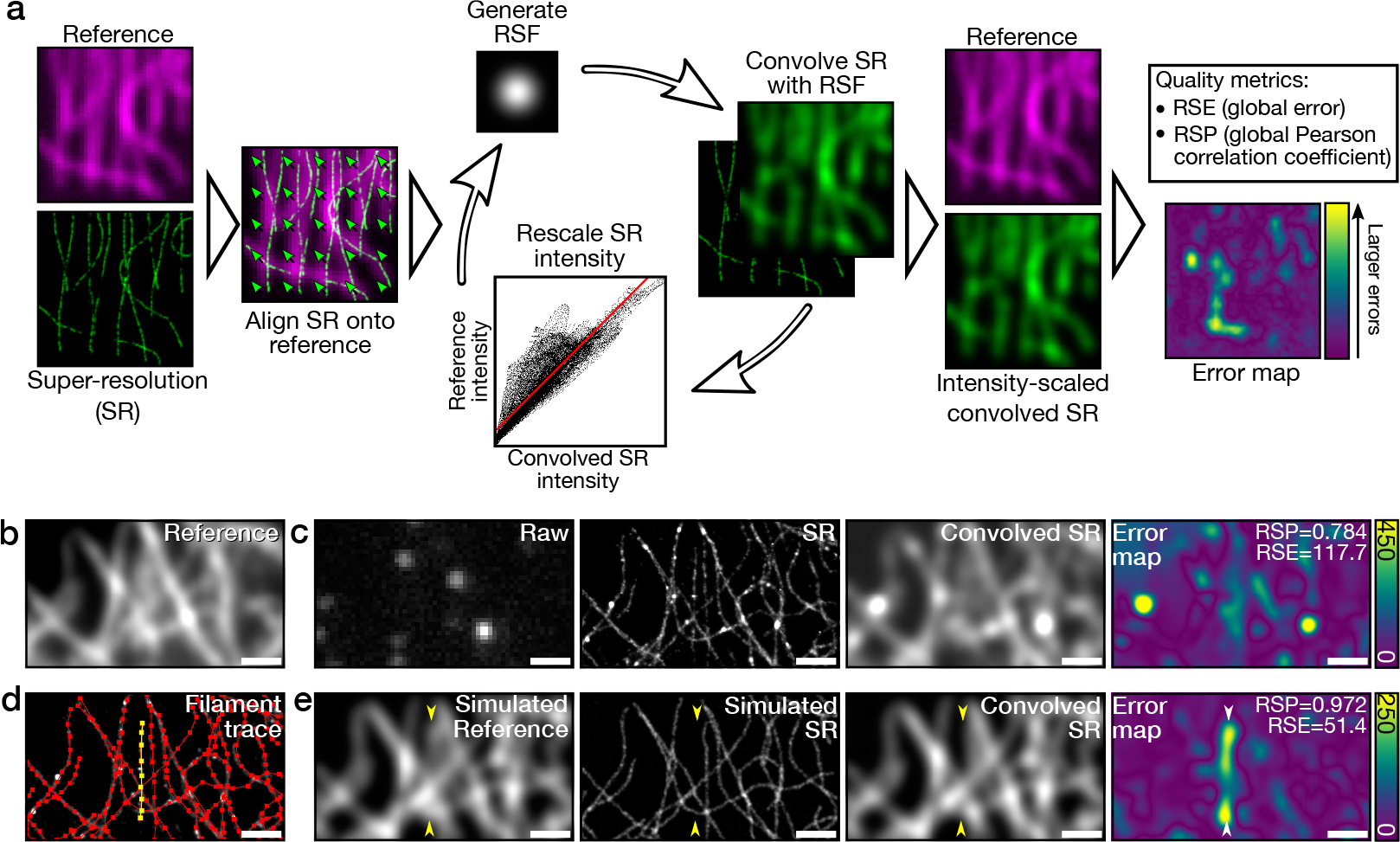
Overview of quantitative error mapping with SQUIRREL. a) Representative workflow for SQUIRREL error mapping. b) Fixed microtubules labelled with Alexa Fluor 647 imaged in TIRF. c) Raw - single frame from raw dSTORM acquisition of structure in **b**, SR - super-resolution reconstruction of dSTORM data set, Convolved SR - super-resolution image convolved with appropriate RSF, Error map - quantitative map of errors between the reference and convolved SR images. d) SuReSim (17) filament tracing used to generate **e**, yellow filament is made to be present in reference image but absent in super-resolution image. e) Simulated reference image, super-resolution image, and super-resolution image convolved with RSF and error map. Yellow arrowheads indicate position of yellow filament seen in **d**. Scale bars = 1 μm.

## Results

**SQUIRREL algorithm.** The SQUIRREL method (Superresolution Quantitative Image Rating and Reporting of Error Locations), implemented as the NanoJ-SQUIRREL plugin, is provided as part of the NanoJ high-performance superresolution data analysis package. It takes advantage of analytical features associated with NanoJ-SRRF (16) and NanoJ-VirusMapper (20). SQUIRREL operates on the assumption that all super-resolution images are representations of fluorescently labelled biological structures rendered at subdiffraction limit resolutions and signal intensities proportional to the local sample labelling density. Thus, by inducing artificial resolution loss in a super-resolution image SQUIRREL generates a new image with a high degree of similarity to an equivalent diffraction-limited one. Quantitative comparison between these two images results in the generation of an error map highlighting regions of high dissimilarity. Such regions point out potential defects in the super-resolution image. The algorithm requires three inputs: a reference image (generally diffraction-limited), a super-resolution image, and a representative resolution scaling function (RSF) image. The RSF can be estimated through optimisation, provided by the user, or for images where the resolution is a ∼10-fold improvement on the diffraction limit, approximated to the PSF of the microscope (Sup. Note 1). Importantly, the RSF should be chosen such that convolution of the superresolution image with the RSF maximizes similarity to the reference image. The process of error mapping starts by the calculation of *I*_RS_, the image created by applying the RSF to the super-resolution image.

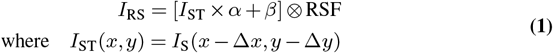

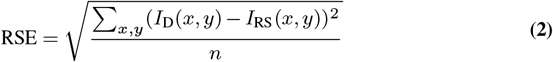

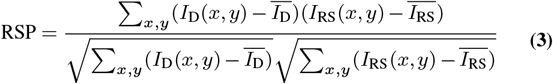

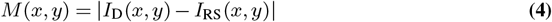

Here, *I*_RS_ is calculated by convolving *I*_ST_ (the result of applying a constant translation correction Δ*x*, Δ*y* to *I*_S_), with the RSF following linear intensity rescaling of *I*_ST_ by constants *α* and *β* (eq. 1, Fig. 1a). The translation is needed to correct for aberrant shifts in the super-resolution image *I*_S_ arising from uncorrected sample drift, differences between the optical path used to collect the reference diffraction-limited image *I*_D_ and *I*_S_, or offsets introduced by reconstruction algorithms. *α* and *β* are required to help match the intensity range of *I*_RS_ with that of *I*_D_. The global similarity between *I*RS and the reference diffraction-limited image *I*D can be calculated through a root-mean-square-error (eq. 2), named RSE for Resolution Scaled Error, and a Pearson correlation coefficient, named RSP for Resolution Scaled Pearson coefficient (eq. 3). Here *n* represents the total number of pixels (where necessary, the dimensions of *I*_RS_ are scaled to match those of *I*_D_), 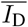 the average value of *I*_D_ and, 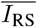 the average value of *I*_RS_. The constants Δ*x*, Δ*y*,α,β can be estimated by optimisation as described in Sup. Note 2. An error map *M* can be generated by calculating the pixel-wise absolute difference between *I*_D_ and *I*_RS_ (eq. 4, Fig. 1a). The root-mean-square-error represented by RSE and Pearson correlation coefficient represented by RSP are adaptedfrom simple metrics classically used to evaluate data similarity, and provide complementary information for assessing image quality. The RSE represents the intensity distance between both images; this measurement is more sensitive to differences in contrast and brightness than the RSP, where the intensity distances are normalised by the mean of each image. The RSP is based on a normalised correlation and its value is truncated between −1 and 1. It thus provides a global score of image quality which can be compared across different super-resolution modalities. While other metrics could be adapted for super-resolution images, such as the structural similarity index (SSIM) (21), here we show that the RSE, RSP and error map provide a robust, detailed evaluation of super-resolution image quality suitable for optimising superresolution experiments. Notwithstanding, both the reference and super-resolution images need to represent the same focal volume. Sup. Note 3 explores how out-of-focus information affects SQUIRREL metrics. It highlights that widefield based references on thick samples compromise the metrics’ fidelity, while this effect will be minimal in optical-sectioning systems such as total internal reflection fluorescence (TIRF), confocal or lattice-light sheet microscopes.

**Validation with real and simulated data.** To demonstrate the capacity of SQUIRREL to identify defects in a superresolution image, we have collected diffraction-limited total internal reflection fluorescence microscopy (TIRF) images of Alexa Fluor 647 immunolabelled microtubules (Fig. 1b) and a corresponding direct STORM (dSTORM) (22) dataset. From these we produced an error map indicating areas of high dissimilarity (Fig. 1c). Regions surrounding filament junctions and overlapping filaments, where the increased local density of fluorophores limits the capacity for singlemolecule localisation, were particularly dissimilar. Based on this, we generated two simulated dSTORM datasets using the SuReSim software (17) (Fig. 1d): a simulated optically realistic reference dataset containing all the traced filaments, and a structural artefact dataset in which a filament was removed. Comparison of the associated reference diffraction-limited images produced an error map that clearly highlights the absence of the selected filament (Fig. 1d). Equally, Fig. S3 shows a similar effect on a SIM acquisition. This result exemplifies the power of SQUIRREL to identify large scale image artefacts in instances where subjective comparison of the widefield (i.e. Simulated Reference) and super-resolution (i.e. Simulated SR) images would be insufficient. To define the smallest scale on which SQUIRREL can identify artefacts, we carried out simulations of an 8 molecule ring structure of varying size. Here, SQUIRREL shows the capacity to quantify image anomalies as small as 100 nm for a noise-free reference, and 150 nm for typical signal-to-noise ratios (Sup. Note 4).

**Comparison between image quality and resolution.** While image resolution is commonly used as a reporter of image quality, in the case of super-resolution studies these two factors weakly correlate (11, 12, 23). One of the favoured methods used in super-resolution and electron microscopy to estimate global resolution across an entire image is Fourier Ring Correlation (FRC) (23). For FRC mapping we assembled and packaged an algorithm within NanoJ-SQUIRREL capable of forming a FRC resolution map of an image by block-wise analysis (Sup. Fig. S5). In Figure 2 we map the local FRC-estimated resolution of a dSTORM dataset for comparison against the SQUIRREL error map. Highlighting various regions of the dataset (Fig. 2a-b), we show that high FRC resolution does not necessarily associate with low error (Fig. 2c-f). In comparison, SQUIRREL error mapping allows for direct visual detection of structural anomalies without coupling quality to a resolution description. As expected, the accuracy of error maps is limited by the resolution of the reference image, the signal-to-noise ratio of the reference and super-resolution images, and the accuracy of the chosen RSF. Nonetheless, SQUIRREL provides a realistic description of local image quality beyond the capacity of a simple resolution estimate.

**Fig. 2.**
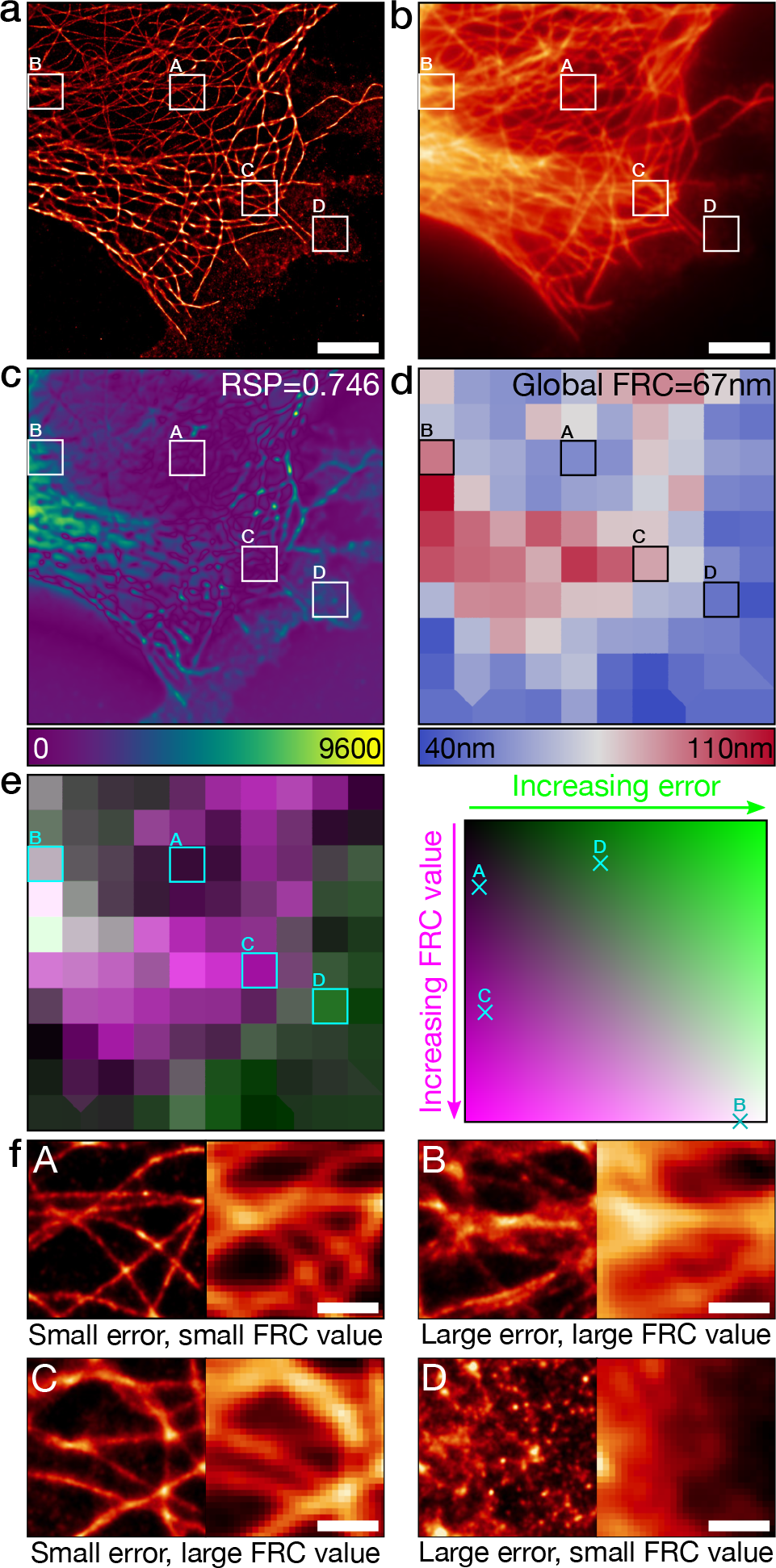
Error mapping and FRC analysis. a) Super-resolution image of fixed Alexa Fluor 647-labelled microtubules reconstructed via MLE. Scale bar = 5 pm. b) Corresponding TIRF image. Scale bar = 5 pm. c) Error map for super-resolution image in **a** using **b** as the reference. d) Local mapping of FRC values for the super-resolution image in **a**. e) Left: Merge of FRC map (magenta) and error map (green, binned to match FRC map). Right: Map of error-resolution space showing where the boxed regions A, B, C and D are located. f) Enlargements of the super-resolution (left) and widefield (right) boxed regions indicated on panels **a**-**e**. Scale bars = 1 μm.

**Super-resolution cross-validation.** The resolution of the error maps produced by SQUIRREL is limited by the resolution of the reference image used. Therefore, in the case of a diffraction-limited reference, the resolution map will only highlight large scale artefacts (Sup. Note 4). It is possible, however, to cross-validate different super-resolution methods using a super-resolution reference. As a pilot sample we chose the prototypic poxvirus, vaccinia virus (VACV). The major infectious form of VACV, mature virions (MVs), are brick shaped particles measuring 360x270x250 nm (24). MVs are composed of three main viral substructures: a biconcave dumbbell shaped core containing the viral genome, two proteinaceous structures termed lateral bodies sitting within the core concavities, and a single lipid bilayer membrane that encompasses these structures (Fig. S6a). We recently described these structures in detail using SIM and STED, mapping a subset of molecular constituents of the virus (20). MVs provide an ideal test case for SQUIRREL as virion substructures cannot be discriminated by conventional fluorescence microscopy but are sufficiently large to be perceived as independent structures by most super-resolution methods. To generate same field-of-view widefield, SIM, STED, maximum likelihood estimation (MLE) and superresolution radial fluctuations (SRRF), where MLE and SRRF images were reconstructed from the same LM dataset, recombinant viruses containing a GFP-tagged lateral body protein (and Alexa Fluor-647-tagged anti-GFP nanobodies for LM imaging) were bound to gridded coverslips and imaged using different optical devices. As various imaging modalities have different optical paths, the acquired images cannot be directly aligned due to optical aberrations. To correct this we have developed a non-rigid registration algorithm, provided in NanoJ-SQUIRREL, which captures the field distortion of each image against a reference (SIM image in this example, Fig. 3a). Using this information a B-spline based local translation is applied to each image, thus generating new images where there is ideal uniform alignment between modalities (Sup. Note 5). In Fig. 3b we show the evolution of error maps for a single virus when using widefield, SIM, STED or SRRF images as a reference. As lateral bodies are considerably smaller than the diffraction limit, when using a widefield reference image, defects could not be perceived, with RSP values approaching 1.0. Using a super-resolution reference immediately highlights dissimilarities between the super-resolution images. These errors reflect asymmetries and inconsistencies along and within lateral bodies; the RSP and RSE also suggest a non-linear intensity scaling across the super-resolution methods. Figure S6b-c shows the RSP and RSE value distributions of 90 viruses analysed from multiple fields-of-view. Interestingly, the range of values in these distributions demonstrates that there is a high degree of variability between super-resolution images generated by different methods. However, by applying SQUIRREL researchers have the potential to filter super-resolution datasets for structures with a high-degree of correlation across various methods prior to further analysis.

**Fig. 3.**
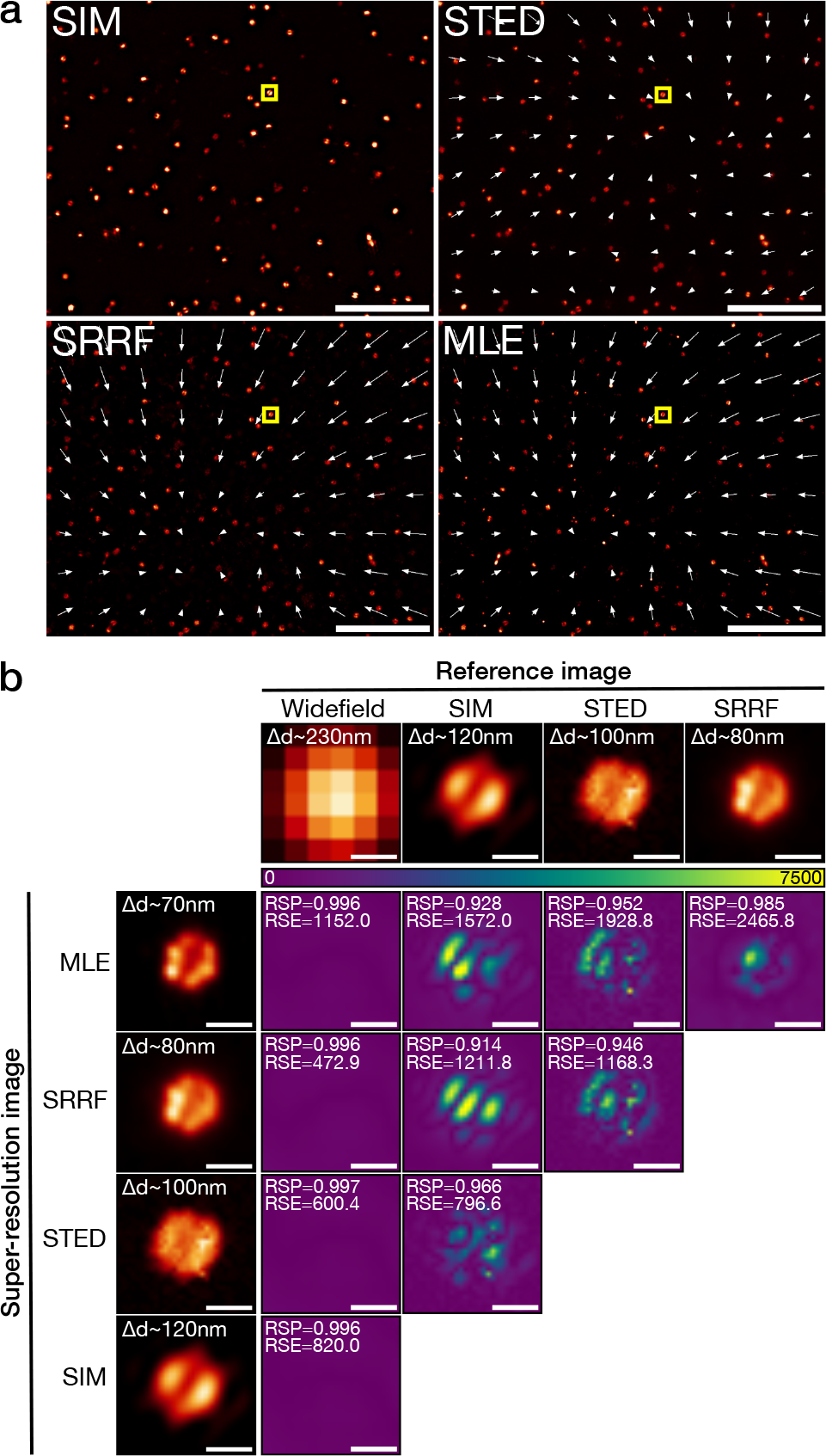
Comparison of vaccinia virus lateral bodies imaged with different super-resolution modalities. a) Field of vaccinia virus particles with GFP-labelled lateral bodies and Alexa Fluor-647 anti-GFP nanobodies imaged with SIM, STED and dSTORM for SRRF and MLE reconstructions. White arrows indicate the transformation used to align images from different modalities with the SIM image. Vector field magnitude was artificially increased 10-fold to aid visualisation. Scale bars = 5 pm. b) Cross-comparisons of the viral particle from the yellow box in **a**. Error maps are displayed for each of the super-resolution modalities when using the widefield images and any super-resolution images with lower resolution as the reference. Scale bars = 200 nm.

**Minimisation of analytical artefacts.** We next asked if SQUIRREL error mapping could be applied to rank, partition and fuse super-resolution reconstructions (Fig. 4a); the potential for such an approach has previously been demonstrated using Richardson-Lucy deconvolution to merge reconstructions with different resolutions (25). To test this a dSTORM dataset of Alexa Fluor 647 immunolabeled micro-tubules was analyzed using three distinct algorithms: ThunderSTORM using a multi-emitter MLE engine (26); SRRF (16); and QuickPALM using a centre-of-mass engine (3). As these methods have the capacity to analyze the same dataset this allowed us to use a single reference diffraction-limited image to rank the reconstructions (Fig. 4c). When doing so we found that, being based on distinct analytical engines, each method yielded a different super-resolution reconstruction where markedly different error distributions could be seen at the same location (Inset Fig. 4b). As the error maps provide spatial details on the local accuracy of each algorithm, they can be converted into weights (Fig. 4d, Sup. Note 6), and the lowest error features of each reconstruction used to generate a new composite image with minimal defects (Fig. 4e). These results demonstrate that image fusion provides an avenue to improve super-resolution by combining information generated using different analytical methods capable of producing images of similar resolution. In addition the RSP and RSE metrics calculated using SQUIRREL enable ranking of images according to their quality, thereby informing researchers on how quality evolves with imaging parameters.

**Fig. 4.**
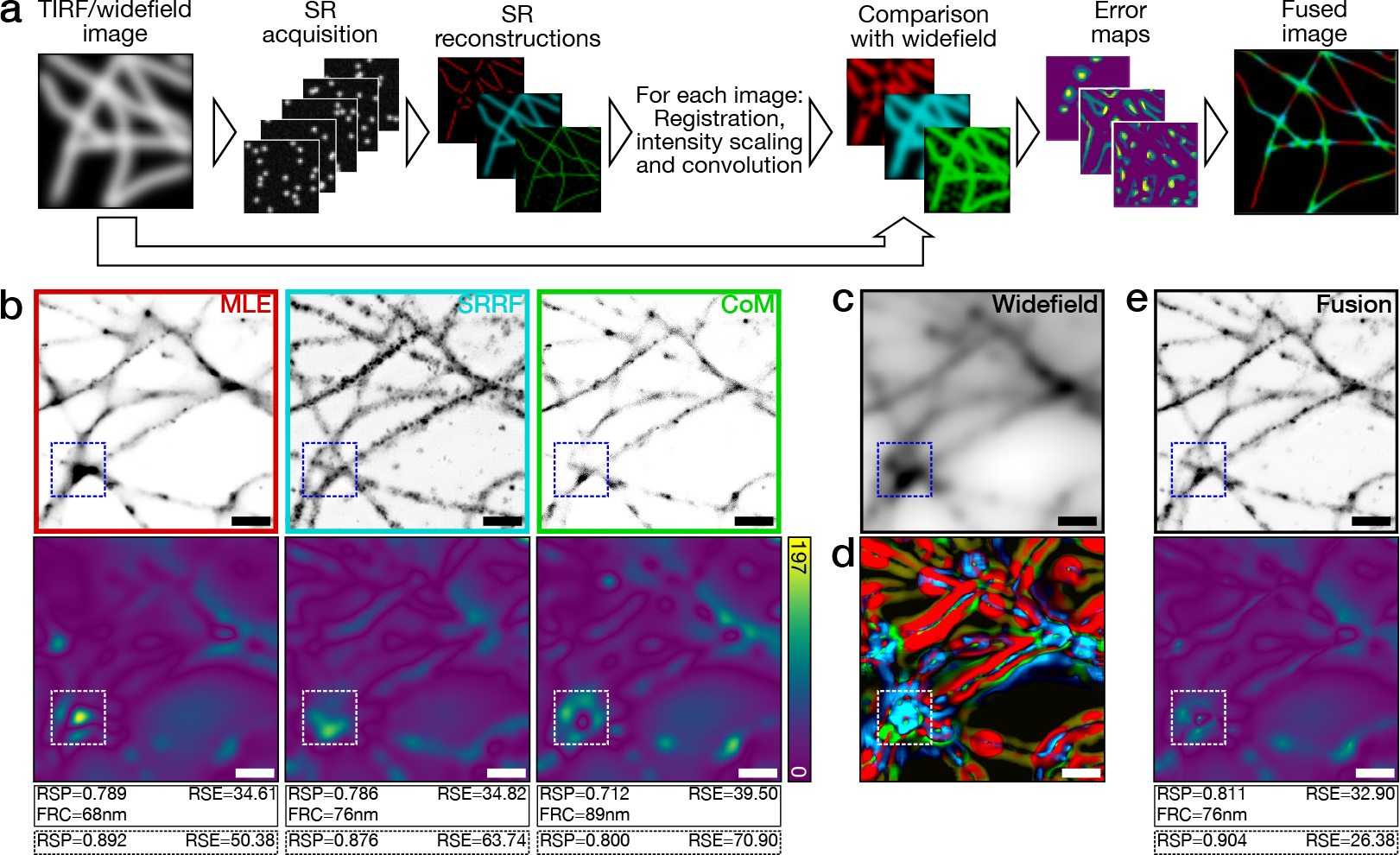
Image fusion of LM data using SQUIRREL. a) Workflow for generating fused images from different super-resolution images from the same PALM/STORM data set. b) Top row: Three super-resolution images generated from the same dSTORM dataset using different algorithms. MLE = maximum likelihood estimatorwith multi-emitter fitting, SRRF = super-resolution radial fluctuations, CoM = centre of mass. Bottom row: Corresponding error maps with the widefield image shown in **c** used as the reference. d)Contributions of different images to the final fused images, colour-coded as in the top row of **b**. e) Top: Fused image, Bottom: Error map of fused image with **c** again used as the reference image. Values in solid line boxes indicate the quality metrics of the whole images, values in dashed boxes represent quality values from highlighted inset region only. Scale bars = 1 μm.

**Improving image acquisition.** Error mapping provides researchers with the means to quantitatively determine how different acquisition parameters may improve the quality of super-resolution images. To test this, a 60,000 frame dSTORM acquisition of sub-diffraction, periodically organized neuronal actin rings was acquired (Fig. 5) (27, 28). Particle localisation was performed, and the localisations used to generate 120 separate super-resolution reconstructions consisting of 500 to 60,000 frames (Fig. 5b). At low frame counts reconstructions showed no periodicity and the error maps exhibited large, widespread errors. As additional frames were included, reconstruction errors decreased and the expected repetitive pattern emerged (Fig. 5d). Surprisingly, the RSP value peaked at 29,000 frames rather than converging at a maximum (Fig. 5e), while inclusion of more than 29,000 frames in the reconstruction led to a small decrease in the RSP value. The same trend was observed for the RSE value, which reached a minimum at 29,000 frames (Fig. 5f). To evaluate if the number of frames used for the superresolution reconstruction had an impact on the visibility of structural details, we quantified the expected periodicity (200 nm) of the actin rings (Fig. 5g). Clearly defined periodicity emerged only when 29,000 or more frames were used for reconstruction (Fig. 5h). Consistent with this, the visibility index of actin rings (Fig. 5i, basis described in Methods and (16)) reached a maximum at 28,500 frames. Addition of more frames to the reconstructed super-resolution image resulted in deterioration of actin ring visibility. The RSP and RSE value peaks, validated by structural analysis, suggest that beyond this maximum quality peak point the unwanted contribution of free label and false detections is greater than the partially depleted correct structural labelling. Fourier analysis (28) of the 29,000 frame reconstruction was in agreement with previously reported values (Fig. 5j). These results indicate that a finite number of frames (29,000) as determined quantitatively by NanoJ-SQUIRREL were optimal for imaging neuronal actin rings, and that any frames acquired beyond this point did not improve image quality. Notably, this information has enormous potential for time-saving during both image acquisition and analysis for any repeatedly performed long acquisition. For the case demonstrated, the acquisition should be done in half the time achieving a higher quality than in a longer acquisition, with this estimation performed quantitatively instead of being based on a subjective human guess. We further performed a similar analysis calculating the evolution of resolution for the same dataset (through FRC calculation, Fig. S7). While FRC analysis indicates improved resolution as the number of frames is increased, the resolution stabilises if 29,000 or more frames are used without any deterioration in resolution beyond this point. Again, as in Fig. 2, this indication that image resolution and image quality are weakly related, and that error mapping provides a more sensitive metric for data reliability.

**Fig. 5.**
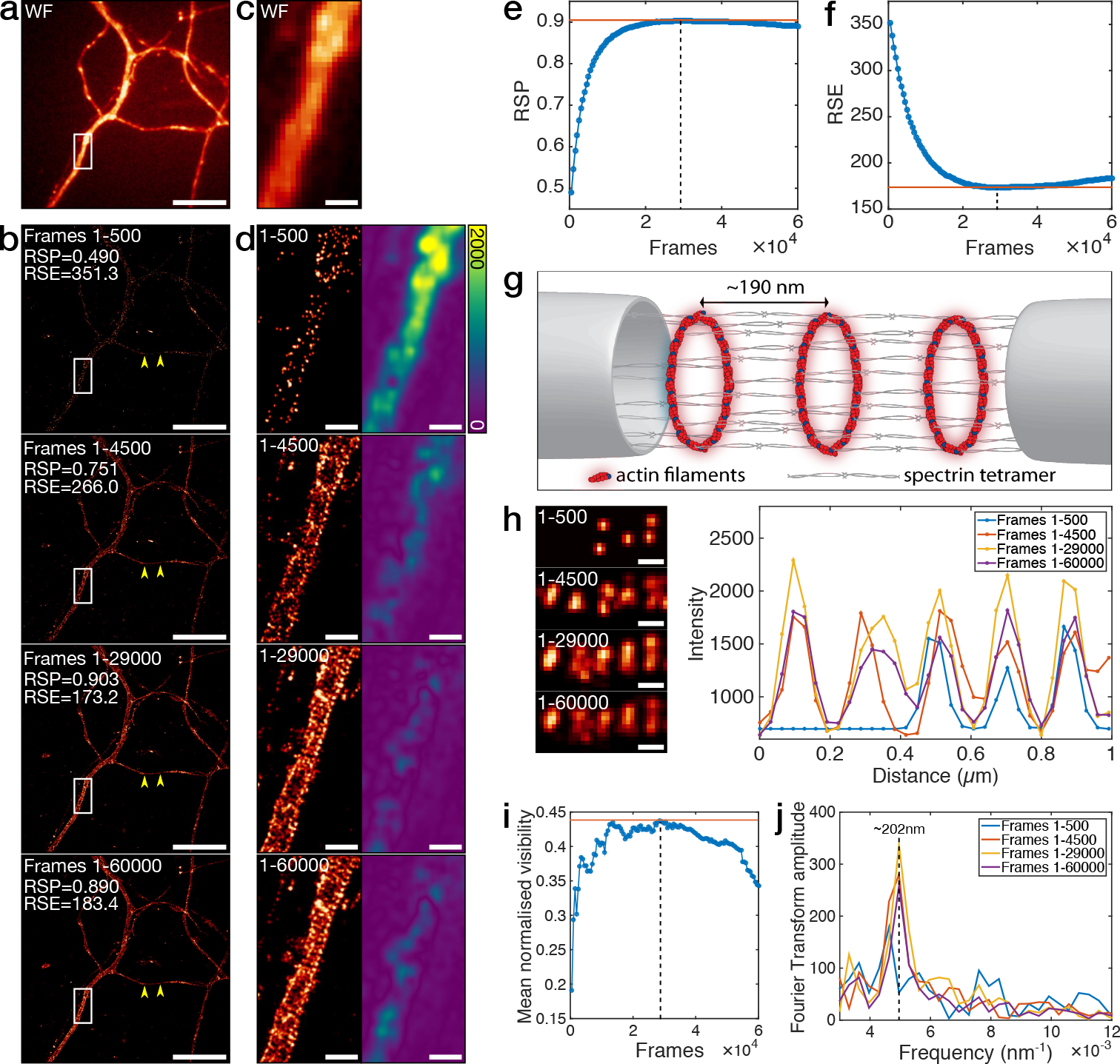
Image quality assessment for informing number of frames for dSTORM imaging of neuronal actin rings. a) Wide-field image of fixed neurons stained with phalloidin-Alexa Fluor 647. Scale bar = 10 pm. b) Super-resolution reconstructions of dSTORM data. The whole raw data set consisted of 60000 frames; four reconstructions are shown with localizations from the first 500 frames, first 4500 frames, first 29000 frames and all 60000 frames. RSP and RSE values were calculated with the wide-field image in **a** acting as the reference. Scale bars = 10 pm. c) Enlarged view of the boxed region in **a**. Scale bar = 1 pm. d) Enlarged views of the boxed regions in **b** and corresponding error maps. Colour bar is consistent for each image. Scale bar = 1pm. e) RSP values for super-resolution images reconstructed from increasing numbers of frames. Dashed line indicates peak RSP value, obtained at 29000 frames. f) As in **e** but for RSE values. Dashed line indicates minimum RSE value, obtained at 29000 frames. g) Schematic showing the arrangement of actin rings and spectrin filaments within an axon. h) Left: 1 pm section of axon displaying periodic actin structures. Scale bars = 200 nm. Right: Line profiles of intensities (scaled to that of the widefield image) along this section of axon. i) Visibility analysis of the line profiles taken along the region in **h** with increasing numbers of frames. Dashed line indicates that the maximum visibility was obtained at 28500 frames. j) Fourier transforms of a 3 pm section of axon as (between the arrowheads in **b**).

**Optimising sample preparation.** Sample labelling is a major element influencing super-resolution imaging quality. LM methods in particular depend on labelling density and fluorophore photoswitching behaviour compatible with the analytical method chosen (10, 16). To ascertain if SQUIRREL image quality readouts can guide optimisation of sample labelling we applied SQUIRREL error analysis to DNA-PAINT based imaging. This method relies on the transient hybridization of complementary DNA templates; the ‘docking’ strand which is appended to the target protein and the fluorescently labelled ‘imager’ strand which is added to the sample media. When combined with LM reconstruction algorithms, it is possible to detect and localise these temporary immobilisation events as a fluorescent spot (29, 30). As bleached fluorophores are not permanently attached to the structure of interest, the on/off kinetics of DNA-PAINT are dictated by the concentration of imager strand utilized (31, 32). Thus we asked if SQUIRREL error analysis could be applied to determine the optimal imager strand concentration for super-resolution imaging of clathrin-coated pits (CCPs). CCPs appear as diffraction-limited spots in wide-field microscopy images (Fig. 6a), and as 100-200 nm rounded pits by super-resolution microscopy (33). Docking strand-labelled clathrin light chain was imaged for 20,000 frames using five different dilutions of imager strand (Fig. 6b). SQUIRREL was used to rank the quality of images produced using three different reconstruction algorithms; MLE, SRRF, and CoM. Displayed in Figure 6c-f are the SQUIRREL-generated error maps for each algorithm at its most and least compatible imager strand concentrations (Fig. 6c-f). Interestingly, the quality of the reconstructed images depended on the combination of imager strand concentration and algorithm used, with the three algorithms showing optimal performance at different imager strand concentrations (Fig. 6h-i). The error maps illustrate the interdependence of sample preparation and algorithm choice as well as how sub-optimal sample preparation leads to increased errors and subsequent disappearance of structures (Fig. 6f, MLE), increased background signal (Fig. 6f, MLE, SRRF) and bridging between structures (Fig. 6f, CoM). Collectively these results demonstrate the utility of SQUIRREL to identify the optimal sample preparation and algorithm combination for a given LM experiment.

**Fig. 6.**
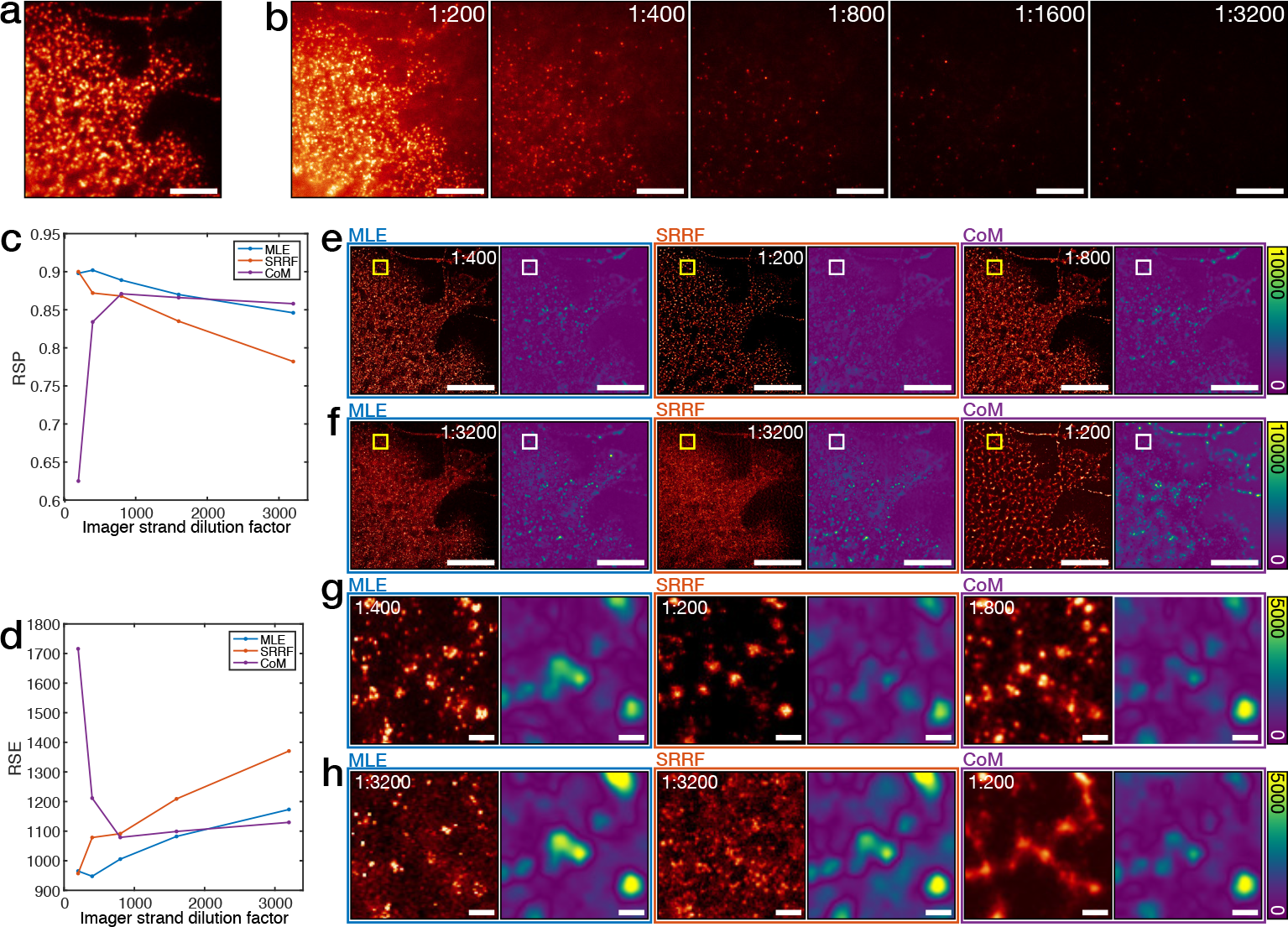
Error analysis for optimisation of DNA PAINT labelling of clathrin-coated pits. a) Widefield image of clathrin-coated pits (CCPs). Scale bar = 5 pm. b) Example frames of the same region as imaged in **a**, but with dilutions of the imager strand at 1:200, 1:400, 1:800, 1:1600 and 1:3200. Scale bars = 5 pm. c) RSP values for MLE, SRRF and CoM reconstructions of the data sets with different imager strand dilutions. d) As in **c**, but RSE values. Super-resolution images and error maps from MLE, SRRF and CoM reconstructions at the best performing imager strand dilution as determined by the RSP values. Scale bars = 5 pm. f) As in **e**, but instead reconstructions are instead from the worst performing imager strand dilution. Scale bars = 5 pm. g) Insets from the boxed regions in **e**. Scale bars = 100 nm. h) Insets from the boxed regions in **f**. Scale bars = 100 nm.

## Discussion

Super-resolution imaging techniques have been established for more than a decade. Yet the fidelity of the data generated is still highly dependent on sample preparation, imaging and data analysis conditions. The complexity of optimising these various parameters without a clear definition of image quality often leads to the erroneous inclusion of image defects. Researchers often benchmark super-resolution images using ‘conventional’ counterparts to validate the preservation of the imaged structures. The NanoJ-SQUIRREL software package provides a quantitative approach to assess superresolution image quality, thereby improving upon subjective visual inspection with error maps and global metrics. We demonstrate that this approach can generate a metric of image quality (Fig. 1a, equations (2) to (4)), uncover the presence of image defects (Fig. 1–Fig. 3), and guide researchers to improved imaging conditions (Fig. 4–Fig. 6). In most publications researchers use a global resolution value as an indicator of image quality. However FRC resolution is often not homogeneous throughout a sample, as seen in Fig. 2. We show that background regions lacking structure frequently have low FRC values which contribute to an overly optimistic estimate of the overall resolution in an image. As an example, the FRC value obtained by averaging only local FRC values in regions of the image containing microtubules in Fig. 2 is 75 nm, compared to the global FRC value of 67 nm. To improve this we have also implemented an algorithm in SQUIRREL to map local resolution values (estimated by FRC) in tandem to error maps. Using this we demonstrate that local resolution does not necessarily correlate with local image quality. Thus, while resolution is important and traditionally used as the hallmark of a good super-resolution image, it should by no means be used as the definition of image quality. The performance of SQUIRREL image error reporting is dependent on the quality of the reference image. While error maps identify discrepancies between the reference and super-resolution images, they cannot indicate which image the error stems from. This is best illustrated when comparing different super-resolution modalities (Fig. 3), in which there is noticeable error conservation when SIM is used as the reference image. This places the responsibility of identifying the source of image errors, or alternatively exclusion of high-error regions from further analysis, on the user. Several publications provide guidelines on optimising super-resolution imaging (4, 5, 7, 10, 34, 35). Importantly, SQUIRREL extends these works by providing a critical quantitative evaluation of how imaging parameters impact upon the fidelity of the final super-resolution image. We show how image quality can be improved at multiple stages of super-resolution imaging, including sample labelling (Fig. 6), choice of imaging modalities (Fig. 3), acquisition settings (Fig. 5) and analysis parameters (Fig. 4). We demonstrate that SQUIRREL provides a quick and easy approach to immediately improve super-resolution data acquisition and quality: from fusion of multiple super-resolution images of the same structure into a new super-resolution estimate of higher quality (Fig. 4b); to informing LM data acquisition to assure optimal throughput and peak quality while preventing oversampling (Fig. 5); and finally, guiding LM labelling conditions such that they are most compatible with the chosen analytical method. Although here we have focused on the application of SQUIRREL to LM images, it can be used with any super-resolution modality provided a reference image is also acquired. Being an open-source ImageJ plugin, NanoJ-SQUIRREL is highly accessible, cross-platform and easily extendible. In the future, we envisage that SQUIRREL will be used for continual monitoring of super-resolution image quality during acquisition. Such a feedback approach could be paired with automated procedures that allow for adaptation of acquisition parameters to assure optimal image quality, reduced acquisition times, and alleviation of data storage requirements.

## Software Availability

NanoJ-SQUIRREL can be downloaded and installed in ImageJ and Fiji automatically by following the instructions in the manual, available here: https://bitbucket.org/rhenriqueslab/nanoj-squirrel.

## ACKNOWLEDGEMENTS

We thank Dr Alex Knight at HolistX Ltd and Dr Seamus Holden at Newcastle University for critical reading of the manuscript. We thank Dr Jonas Ries at European Laboratory for Molecular Biology (EMBL) Heidelberg for provision of customised MATLAB software and critical reading of the manuscript. Kalina Tosheva at University College London (UCL) for critical reading of the manuscript and beta testing the software. We thank Dr Buzz Baum at UCL for reagents. Many of the look-up-tables used here used are based on the open-source repository of David Williamson at King’s College London. This work was funded by grants from the UK Biotechnology and Biological Sciences Research Council (BB/M022374/1; BB/P027431/1; BB/R000697/1) (R.H. and PM.P), core funding to the MRC Laboratory for Molecular Cell Biology at University College London (J.M.), the European Research Council (649101—UbiProPox) (J.M.)., the UK Medical Research Council (MR/K015826/1) (R.H. and J.M.), the Wellcome Trust (203276/Z/16/Z) (S.C. and R.H.) and the Centre Nationnal de la Recherche Scientifique (CNRS ATIP-AVENIR program AO2016) (C.L.). D.A. is presently a Marie Curie fellow (Marie Sklodowska-Curie grant agreement No 750673). C.J. funded by a Commonwealth scholarship, funded by the UK government.

## AUTHOR CONTRIBUTIONS

S.C. and R.H. devised the conceptual framework and derived theoretical results. S.C., D. A., C.L., J.M. and R.H. planned experiments. S.C. and R.H. wrote the algorithm. Simulations were done by S.C. Experimental data sets were acquired by S.C. (Fig. 1b), D.A. (Fig. 2,Fig. 3), C.J. (Fig. 2), P.M.P. (Fig. 4) and C.L. (Fig. 5-Fig. 6). Data analysed by S.C. and D.A. while C.L., J.M. and R.H. provided research advice. The paper was written by S.C., D.A., J.M. and R.H. with editing contributions of all the authors.

## Methods

**Cell lines and primary cells.** HeLa cells were kindly provided by Prof Mark Marsh, MRC LMCB, UCL and cultured in phenol-red free DMEM (Gibco) supplemented with 2 mM GlutaMAX (Gibco), 50 U/ml penicillin, 50 μg/ml streptomycin and 10% fetal bovine serum (FBS; Gibco). CHO cells were cultured in phenol red-free Minimum Essential Medium Alpha (MEMα; Gibco) supplemented with 10% FBS (Gibco) and 1% penicillin/streptomycin (Sigma). Rat hippocampal neurons and glial cells were harvested from embryonic day 18 pups, following established guidelines of the European Animal Care and Use Committee (86/609/CEE) and approval of the local ethics committee (agreement D13-055-8), and cultured in Neurobasal medium (Gibco) supplemented with 2 mM GlutaMAX-I (Gibco) and B27 supplement (Gibco). All cells were grown at 37°C in a 5% CO_2_ humidified incubator.

**Sample preparation for Widefield and TIRF-SMLM imaging of fixed microtubules.** For TIRF-SMLM imaging of microtubules, 13 mm diameter, thickness #1.5 coverslips were washed overnight in 1:1 HCl:methanol and washed 5 times in ddH_2_O and twice in 90% isopropyl alcohol. Coverslips were then incubated overnight in poly-L-lysine (0.01%) (Sigma Aldrich) and rinsed twice in PBS. HeLa cells were seeded on these coverslips and grown overnight in 12-well plates. Cells were fixed with 4% PFA in cytoskeleton buffer (10 mM MES, pH 6.1, 150 mM NaCl, 5 mM EGTA, 5 mM glucose, 5 mM MgCl_2_) for 15 min at 37°C, washed 3x with PBS, then permeabilised with 0.1% Triton X-100 in PBS for 10 min and blocked in 2.5% BSA in PBS for a further 30 min. Samples were then labelled with 2 μg/ml anti-a-tubulin (DM1A mouse monoclonal, Sigma Aldrich) in 2.5% BSA in PBS for 1 hour, followed by 3x washes with PBS and labelling with Alexa Fluor 647-labelled goat anti-mouse secondary antibody (Invitrogen) (2μg/ml in 2.5% BSA in PBS) for 1 hour. Samples were washed 3x with PBS and fixed again in 4% PFA in cytoskeleton buffer for 10 min, before being washed 3x with PBS. Samples were mounted on a parafilm-formed gasket (1) in STORM buffer (150 mM TRIS, pH 8, 10 mM NaCl, 1 % glycerol, 1 % glucose, 1 % BME), sealed with clear nail varnish (Maybelline) and imaged within 3 hours of mounting.

For widefield super-resolution imaging of microtubules for image fusion, CHO cells were seeded on ultra-clean (1) 8 mm diameter thickness #1.5 coverslips (Zeiss) at a density of 0.1 x 10^6^ per 35 mm dish. Fixation was performed with 4% PFA in a modified version of cytoskeleton stabilising buffer (CSB) (5 mM KCl, 0.1 mM NaCl, 4 mM NaHCO_3_, 11 mM Na_2_HPO_4_, 2 mM MgCl_2_, 5 mM PIPES, 2 mM EGTA, pH 6.9) for 15 min at 37°C, followed by washing with the same CSB (without PFA). Additional permeabilization was performed (0.05% Triton X-100 in CSB) for 5 min followed by three washing steps using 0.05% Tween-20 in the modified version of CSB and blocking in 5% BSA (Sigma) for 40 min. Microtubules were stained and submitted to a secondary fixation step as described above. 100 nm TetraSpeck beads (Life Technologies) were added at a dilution of 1:1000 in PBS for 10 min to each coverslip. Coverslips were mounted on clean microscope slides (1) in 100 mM mercaptoethylamine (Sigma) at pH 7.3 and all imaging was performed within 3 hours of mounting.

**Sample preparation and imaging of actin and CCPs in fixed neurons and glial cells.** Rat hippocampal neurons or glial cells (from embryonic day 18 pups) were cultured on 18mm coverslips at a density of 10000 /cm^2^ or 4000 /cm^2^, respectively. After 9 days in culture, samples were fixed using 4% PFA in PEM (80 mM PIPES, 2 mM MgCl_2_, 5 mM EGTA, pH 6.8) for 10 min. Preparation of actin-stained neurons for SMLM was performed similarly to the protocol described in (2), with minor modifications. After blocking, neurons were incubated with a mouse anti-map2 primary antibody (Sigma Aldrich, catalogue #M4403) for 1h30 at RT, then with a Alexa Fluor 488 labelled donkey anti-mouse secondary antibody (Thermo Fisher) for 45 min at RT, then with 0.5 mM phalloidin-Alexa Fluor 647 (Thermo-Fisher) overnight at 4 °C. Neurons were mounted in a modified STORM buffer (50 mM Tris, pH 8,10 mM NaCl, 10% glucose, 100 mM mercaptoethylamine, 3.5 U/ml pyranose oxidase, 40 μg/mL catalase) complemented with 0.05 mM phalloidin-Alexa Fluor 647, to mitigate phalloidin unbinding during acquisition and imaged immediately.

For PAINT imaging (3) of CCPs in glial cells, fixed neuron samples were incubated with a rabbit anti-clathrin primary antibody (abCam, catalogue #21679) overnight at 4 °C, then with an anti-rabbit DNA-conjugated secondary antibody (Ultivue) for 1 hour at room temperature.

**VACV sample preparation.** 2.5 ×10^6^VACV particles (WR strain, EGFP-F18 in tk locus (4)) were diluted in 100 μl 1 mM TRIS, pH 8, sonicated for 3x 30s and incubated on gridded #1.5 glass-bottom petri dishes (Zell-Kontakt GmbH) for 1 hour at room temperature and fixed with 4 % PFA in PBS for 15 min. Samples were quenched with 50 mM NH_4_Cl in PBS for 10 min, washed in PBS, and incubated in permeabilization/blocking buffer (1% Triton X-100, 5% BSA, 1 % FBS in PBS) for 30 min. Samples were labelled in blocking/permeabilisation buffer overnight at 4 °C or 2 hours at room temperature with anti-GFP nanobodies (Chromotek), labelled in-house with Alexa Fluor 647 NHS-ester (Life Technologies) with a dye-to-protein ratio of approximately 1, as previously described (5). Samples were then washed 3x with PBS, fixed in 4% PFA in PBS for 10 min, quenched with 50 mM NH_4_Cl in PBS for 10 min and washed in PBS.

**Imaging.** Fixed microtubule samples were imaged by TIRF-SMLM on a N-STORM microscope (Nikon Instruments), using a 100× TIRF objective (Plan-APOCHROMAT 100×/1.49 Oil, Nikon) with additional 1.5× magnification. A reference TIRF image was acquired with 5% power 647 nm laser illumination and 100 ms exposure time, before SMLM data acquisition of 40 000 frames at 100% power 647 nm illumination with 405 nm stimulation and an exposure time of 30 ms per frame.

Widefield and super-resolution imaging of fixed microtubules for fusion was carried out on a Zeiss Elyra PS.1 inverted microscope system, using a 100× TIRF objective (Plan-APOCHROMAT 100×/1.46 Oil, Zeiss) and additional 1.6× magnification. The sample was illuminated with a 642 nm laser operating at 100% laser power. 45000 frames were acquired with a 20ms exposure time per frame.

Neuron samples were imaged on a N-STORM microscope using a 100× objective (Plan-APOCHROMAT 100×/1.49 Oil, Nikon). The sample was illuminated at 100% laser power at 647 nm. A sequence of 60,000 images at 67 Hz was acquired.

DNA-PAINT imaging of CCPs in glial cells was performed on a N-STORM microscope using a 100× objective as above. The same glial cell (present in low numbers in hippocampal cultures) was imaged in serial dilutions of Imager-650 (2 mM stock, from lower to higher concentration) in Imaging Buffer (Ultivue). The sample was illuminated at 647 nm (50% laser power) and a sequence of 20,000 images at 33 Hz was acquired for each Imager-650 dilution, before switching to a higher concentration of Imager-650 in Imaging Buffer.

VACV samples were imaged in STORM buffer on a Zeiss Elyra PS.1 system, using a 100× TIRF objective with additional 1.6× magnification (as above) for SIM, SRRF and SMLM acquisition. Buffer was exchanged to PBS and STED images were acquired on a Leica SP8, re-localising the same ROI based on the grid. SMLM data acquisition parameters were 30 000 frames at 100% laser power 647 nm illumination with 405 nm stimulation and an exposure time of 33 ms per frame.

**Reconstruction algorithms for dSTORM data.** The freely available software packages ThunderSTORM (6), SRRF (7) and QuickPALM (8) were used for superresolution image reconstruction in Figs. 2, 3, 4 and 6. Images labelled ‘MLE’ were reconstructed with ThunderSTORM with the integrated PSF method with maximum likelihood fitting and multi-emitter fitting enabled. Drift correction was performed post-localization and images were rendered using a normalized 20 nm Gaussian. Images labelled ‘SRRF’ were analysed with the most appropriate parameters for each individual data set and drift corrected during analysis. Images labelled ‘CoM’ were reconstructed using QuickPALM with the default parameters, following drift correction of the raw data using the NanoJ-SRRF package. The particle tables from QuickPALM were then loaded into ThunderSTORM for rendering using a normalized 20 nm Gaussian. Images in Fig. 5 were rendered with ThunderSTORM using a normalized 20 nm Gaussian from particle tables generated with SMAP, a MATLAB based software package developed by the Ries group at the EMBL, Heidelberg. Localizations were determined using a probability based method after background subtraction by wavelet filtering and lateral drift was corrected by cross-correlation.

**Super-resolution image simulation with SuReSim.** In order to simulate disappearance of a filament from a realistic microtubule network, a real super-resolution image of microtubules (Fig. 1c) was used as a support for SuReSim data simulation. Raw data of blinking Alexa 647-labelled microtubules imaged using TIRF were reconstructed using ThunderSTORM maximum likelihood multi-emitter fitting and then loaded into the SuReSim software and all filaments were traced using the editor function and the WIMP file saved. SuReSim was used to generate a simulated super-resolution reconstruction of all filaments, which was then convolved by a Gaussian PSF to generate a simulated reference image. The object in the WIMP file corresponding the to filament highlighted in Fig. 1d-e was deleted, and SuReSim was used again to render a simulated super-resolution reconstruction, except this time missing a filament.

**Visibility analysis.** To quantify the quality of the superresolution reconstructions of parallel actin rings, a normalized visibility similar to that described in Geissbuehler et al. (9) was calculated as follows. Average intensity profiles were plotted for a 0.5 x 1 pm stretch of axon containing 5 actin rings (region shown in Fig. 5h) for each of the 120 reconstructed images. The MATLAB function findpeaks was used to find the 5 peak positions in the average profile measured from the 60000 frames reconstruction, and mean pairwise visibility was calculated as follows.

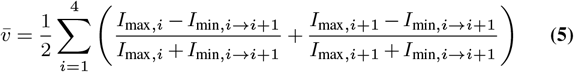

*I*_max_,*i* and *I*_max,*i*+1_ are the intensities at peak positions *i* and *i +1* respectively, where *i* denotes the index of the actin ring in the sampled region, and *I*_min,*i*→*i*+1_ is the intensity at the midpoint of two adjacent peaks. Higher visibilities correspond to a greater ability to differentiate between two structures up to a maximum value of 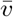 = 0.5.

**Colour maps.** Colour maps used for displaying images (‘NanoJ-Orange’), error maps (‘SQUIRREL-errors’) and FRC maps (‘SQUIRREL-FRC’) are provided in the NanoJ-SQUIRREL software package.

## Supplementary Note 1: Extracting the Resolution Scaling Function

The Resolution Scaling Function (RSF) can be estimated in 3 different ways: A) automatically during the Error-Map generation procedure; B) approximated to the Point-Spread-Function of the microscope; C) derived when the reference PSF and superresolution PSF is known. These processes are described below:

A. Automatic RSF generation.For the majority of cases, a symmetric normalized Gaussian function *(G(x,y))* (Eq. S1) provides a good numerical approximation to the RSF. Studies in microscopy have shown that the PSFs of widefield, Total Internal Reflection Fluorescence Microscopy (TIRF), Laser Scanning Confocal Microscopy (LSM) and Spinning Disk Microscopy (SD) can be, in a simplified manner, approximated to Gaussian functions (1, 2, 3). Similarly, in super-resolution methods, the final images generally present a Gaussian-like PSF (2, 3, 4, 5). The RSF in itself is expected to be a function that by convolution would convert the super-resolution PSF (PSFs) into the reference image PSF (PSF_D_) (Eq. S2); if both of these are assumed to follow a Gaussian distribution, then the RSF will be well approximated by a Gaussian. However, the *σ* value describing the Gaussian RSF is unknown but can be estimated during optimisation. Sup. Note 2 provides an analytical description of this process.

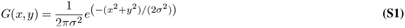

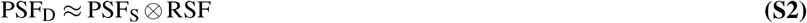

**B. Approximation of RSF to the PSF.** When the resolution of the super-resolution dataset is considerably high (< 30 nm, ∼10-fold improvement upon the diffraction limit), the Point-Spread-Function (PSF) of the microscope provides a good approximation of the RSF. In such cases, the user will be able to provide a PSF rendering to the Error-Map calculation procedure, as a proxy to the RSF. For simplicity, we provide as part of NanoJ-SQUIRREL an algorithm capable of extracting a model PSF from an image composed of sparse sub-diffraction PSF-like particles (Fig. S1). Alternatively, the user can also derive a theoretical PSF based on a diffraction model; the PSF Generator ImageJ-plugin (6) is an excellent tool in such cases, providing PSF rendering capability and maintaining an extensive library of PSF models.

**Fig. S1.**
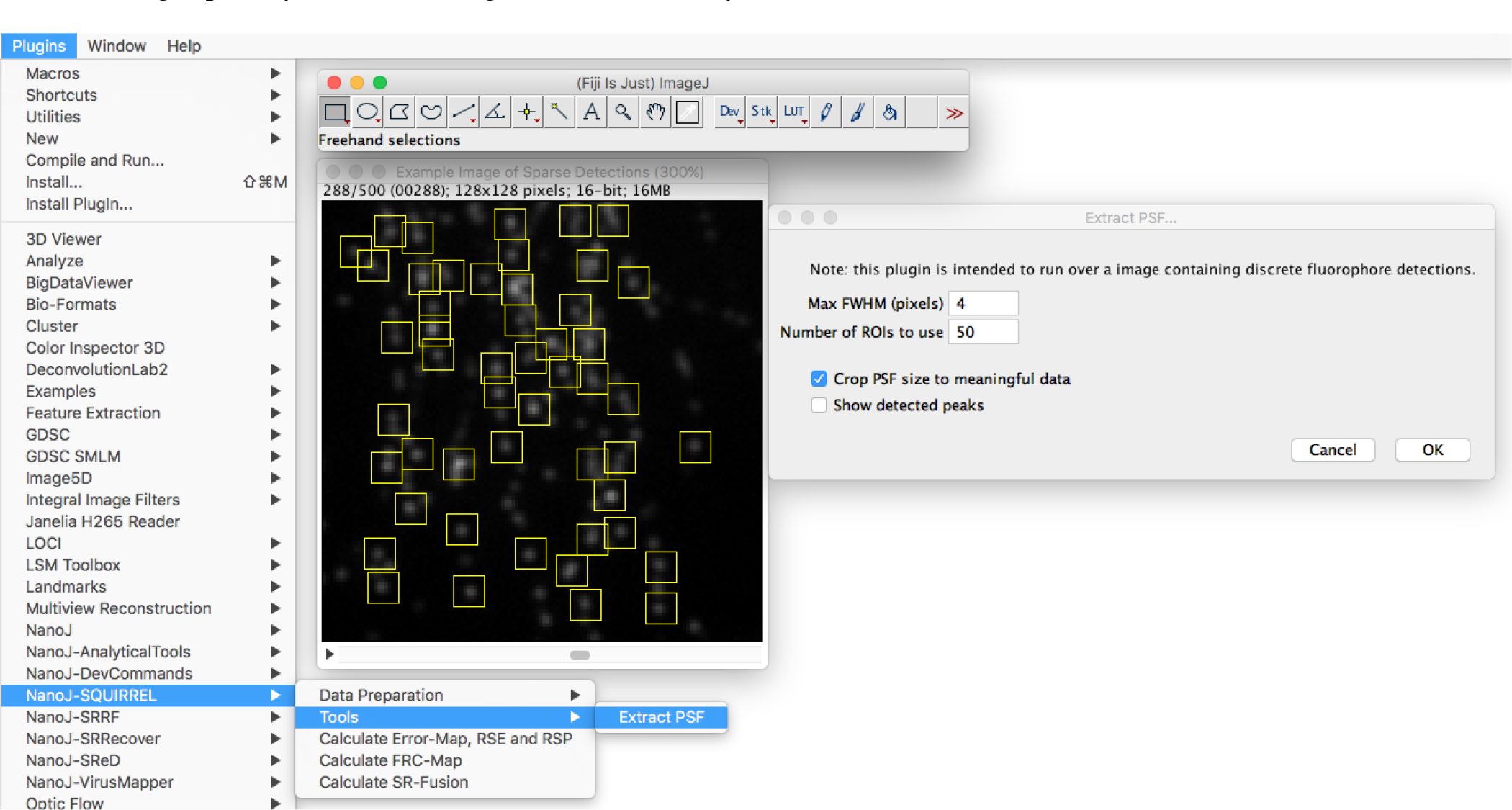
The extract PSF algorithm, part of NanoJ-SQUIRREL. This method will search for peaks in the image (local maxima) and fit a 2D Gaussian function to each (of varying amplitude, centre, background and σ). This fitting estimates the centre of each peak with a sub-pixel accuracy. Each peak is then extracted, aligned and re-rendered in a list of images representing all the detected peaks. An initial estimate is generated by an average projection of all extracted peaks. A final estimate is then regenerated by a weighted average projection where each peak component is weighted by its Pearson correlation coefficient against the previous average projection.

**C. Derived when the reference PSF and super-resolution PSF are known.** Through Eq. S2, the RSF matrix can be estimated if both PSF_D_ and PSF_S_ are known. These two last matrices can be extracted either from real data, or estimated by underlying analytical models. For a noise free case, the calculation of the RSF can be done in Fourier space, by calculating the inverse transform of the division between the Fourier transformed PSF_D_ and Fourier transformed PSF_S_.

## Supplementary Note 2: Estimating the Error Map, RSE and RSP

The process of estimating an Error Map via SQUIRREL is divided in 3 subsequent steps represented in Fig. 1a and described below. The following notation will be used to denote the different images during this process:

- *I*_D_: reference image
- RSF: resolution scaling function
- *I*_RSF_: resolution scaling function integrated over finite pixels
- *I*_S_: original super-resolution image
- *I*_ST_ = *I*_s_ (*x* — Δ*x*, *y* — Δ*y*): super-resolution image registered to reference image
- *I*_STγ_ = *I*_ST_ × α + β: registered super-resolution image following linear intensity rescaling

**A. Registration of Super-Resolution Reconstructions against the Reference image.** The first step of registration is the estimation of Δ*x*, Δ*y* (Eq. 1), through cross-correlating the reference and the super-resolution images. For this purpose, the cross-correlation is calculated through a fast Hartley transform (FHT), taking advantage of the threaded Parallel Colt library (7). Δ*x*, Δ*y* can then be estimated by calculating the spatial difference between the coordinates of the correlation matrix peak intensity value and its geometric centre. To detect the coordinates of the peak correlation value, we selectively upscale the correlation matrix via bi-cubic spline interpolation (8) and find its maximum. This process has been previously described in (9). Finally, we employ a bi-cubic spline translation of the super-resolution image to maximise its overlap to the reference image.

**B. Image intensity rescaling and RSF estimation.** The intention of this step is to linearly rescale the intensity of the superresolution estimate *I*_S_ and convolve it with *I*_RSF_ in a manner that will maximise its similarity to the reference image *I*_D_. To do so, the unknown variables *a* and *3* defining the intensity rescaling shown in Eq. 1 need to be estimated. Additionally, if an RSF is not input, the SQUIRREL algorithm will automatically estimate its matrix by an approximation to a 2D Gaussian function (Eq. S3) of unknown *σ*. Similarly to (10), we integrate the Gaussian function over finite pixels (Eq. S4-S5). The estimation of these two (if RSF is known) or three variables (if unknown) is achieved in SQUIRREL through a highly threaded implementation of a Particle Swarm Optimiser (PSO) (11,12). PSO optimisation is a derivative-free, metaheuristic optimisation approach taking few assumptions about the optimisation problem posed and is capable of searching a large space of candidate solutions. Eq. S6 defines the least-squares minimisation, where the result of *I*_STγ_(α, β) ⊗ *I*_RSF_(σ) is scaled down to the same width and height as *I*D by pixel averaging prior to their subtraction.

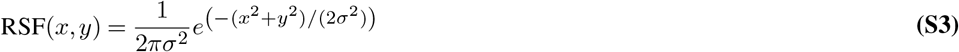

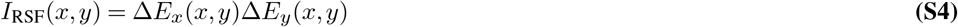

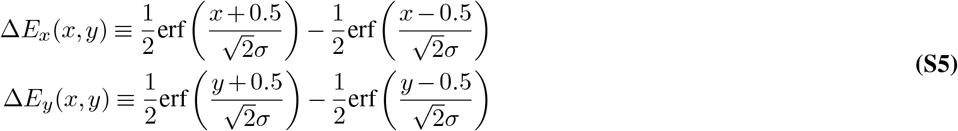

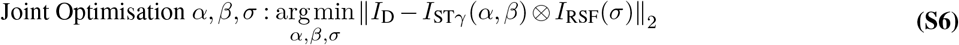

**C. Calculation of error map, RSE and RSP.** Given that α, *β* and RSF are now known, the Resolution Scaled Error (RSE) can be calculated through Eq. 2, the Resolution Scaled Pearson coefficient (RSP) by Eq. 3 and the error map by equation 4.

## Supplementary Note 3: Out-of-focus effects on the Error Map

When using microscopes with poor optical sectioning, for example widefield systems, discrepancies between the reference and super-resolution images can arise from out-of-focus structures. To assess the impact of this effect, a structure was simulated at a linear tilt to the focal plane such that one end of the structure was 750 nm below the focal plane, the centre was in focus, and the other end was 750 nm above the focal plane. SQUIRREL analysis was performed (Fig. S2). Firstly, a ‘defect-free’ single molecule data set was produced by simulating three-dimensional blinking molecules using a Born and Wolf optical model (6). This image sequence was then analysed using ThunderSTORM (13) to produce a realistic super-resolution image and then compared with a simulated widefield image of the same structure (Fig. S2a, c). The error map shows that the superresolution reconstruction was, as expected, accurate for the in-focus regions of the structure in the middle of the image but in poor agreement with the out-of-focus regions of the sample. Next we investigated whether errors arising from out-of-focus fluorescence would mask errors within the focal plane. To do this, a deliberately artefactual single molecule data set was produced where periodic regions of the structure were absent; this was again reconstructed using ThunderSTORM analysis (Fig. S2b). The error map generated through comparison with the defect-free widefield image (Fig. S2c) clearly demonstrates that while the errors arising from out-of-focus fluorescence are still present, the errors resulting from structural mismatch are also clearly identifiable as periodic peaks in the error map.

**Fig. S2.**
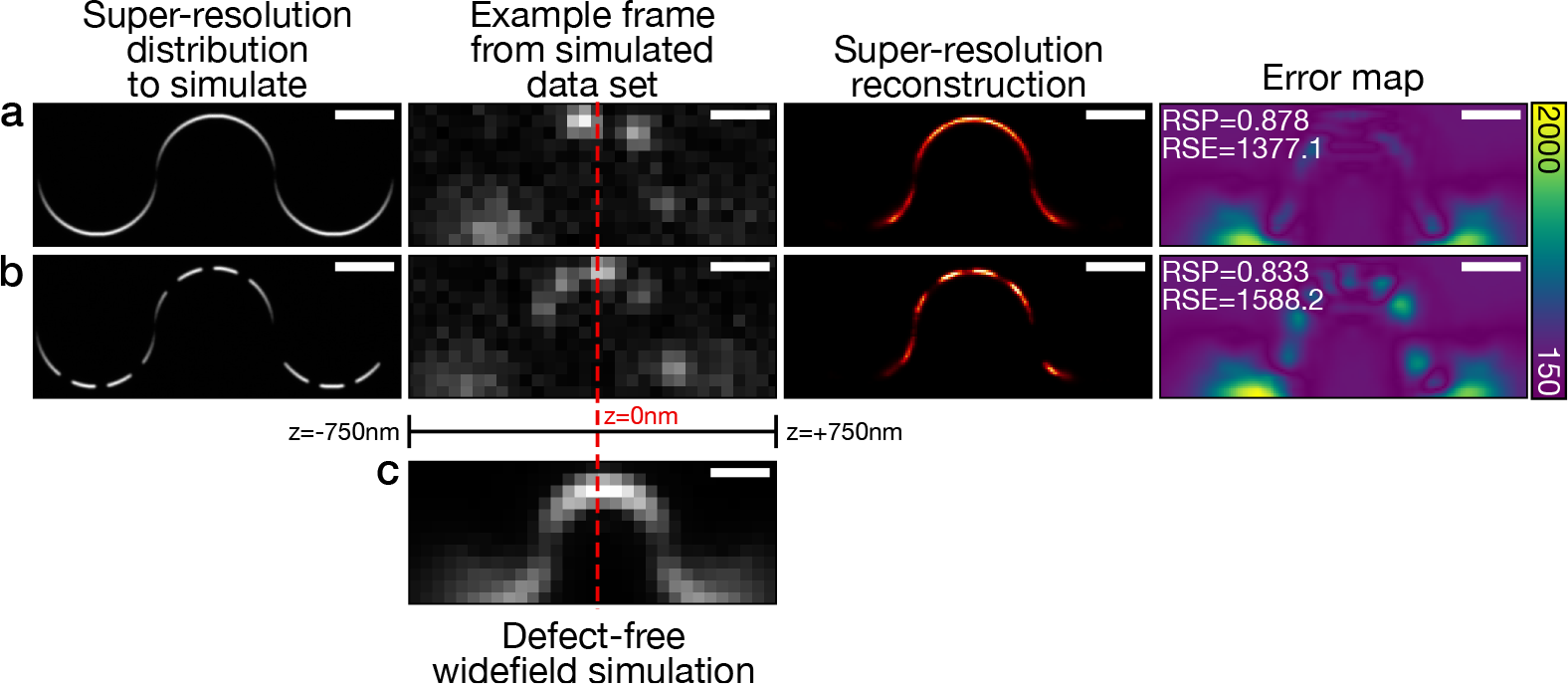
Impact of out-of-focus fluorescence on error maps, RSP and RSE values. a) Test structure for defect-free super-resolution simulation; example frame from simulated single molecule data set where there is an axial tilt along the x-dimension; result of running the simulated data set through ThunderSTORM analysis; error map generated between the reconstructed image and a simulated defect-free widefield image (c). b) As in a, except here there are periodic defects in the super-resolution image which are not present in the widefield image. c) Defect-free widefield image produced of the tilted structure shown in (a). Scale bars 500 nm.

**Fig. S3.**
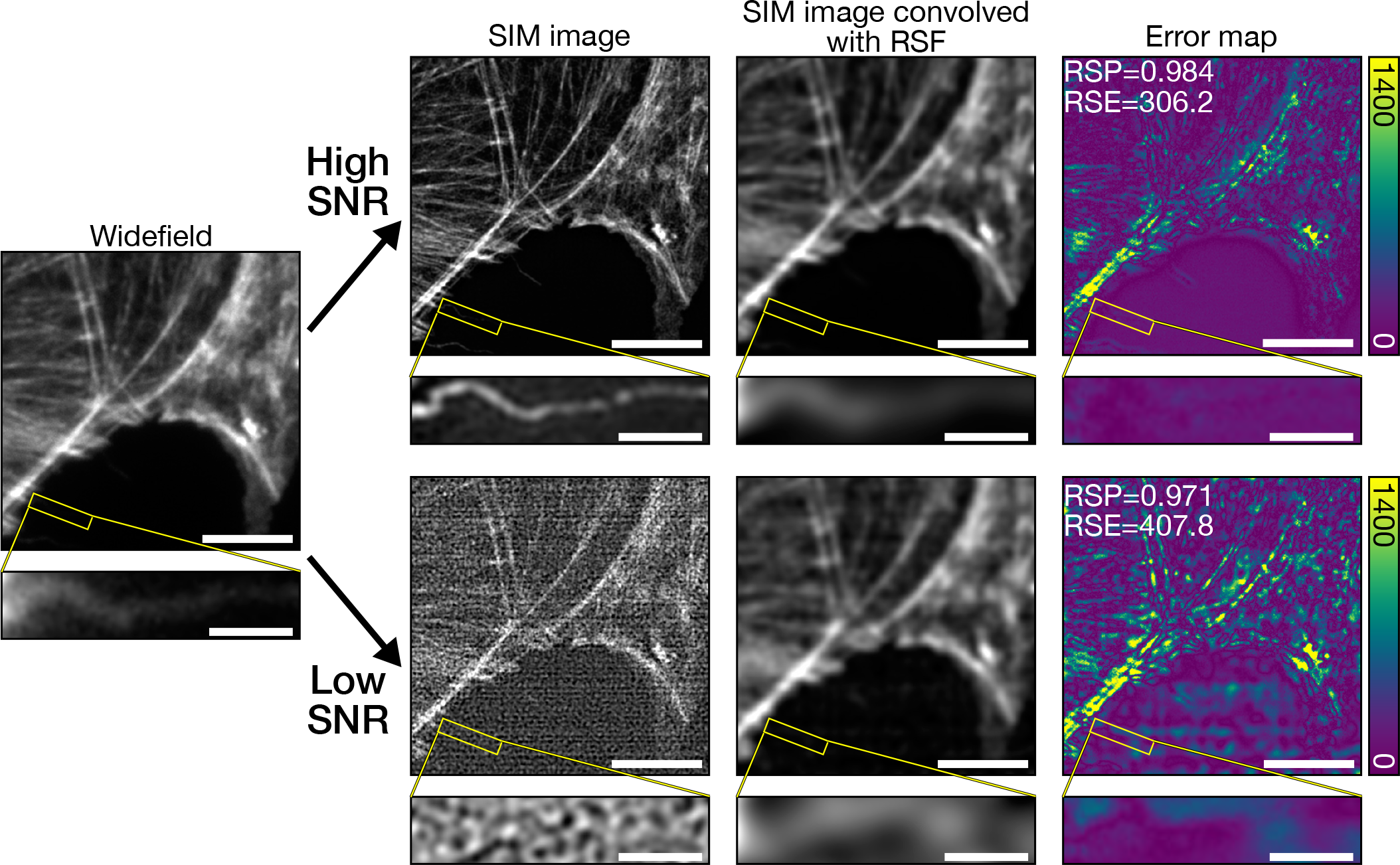
Application of SQUIRREL analysis to SIM data. Actin imaged using standard widefield microscopy followed by SIM at two different laser intensities. SQUIRREL analysis was run on the two SIM images, both using the widefield image as the reference. With an increased laser intensity a high SNR is achieved (top row), yielding more structural detail in the SIM image than with a decreased laser intensity (bottom row). Fine details are lost (inset) and artefactual background patterning (error map) becomes apparent at low SNR. Main image scale bars = 5 μm, inset scale bars = 1 μm.

## Supplementary Note 4: Size and Signal-to-Noise sensitivity of SQUIRREL

To estimate the resolution scale to which SQUIRREL is able to identify super-resolution anomalies, we here designed a simulated super-resolution imaging experiment where controlled defects can be incorporated into a known structure. To this effect, we have simulated an 8-molecule ring structure where the separation between adjacent molecules (Δd) can be varied (Fig. S4a).

**Fig. S4.**
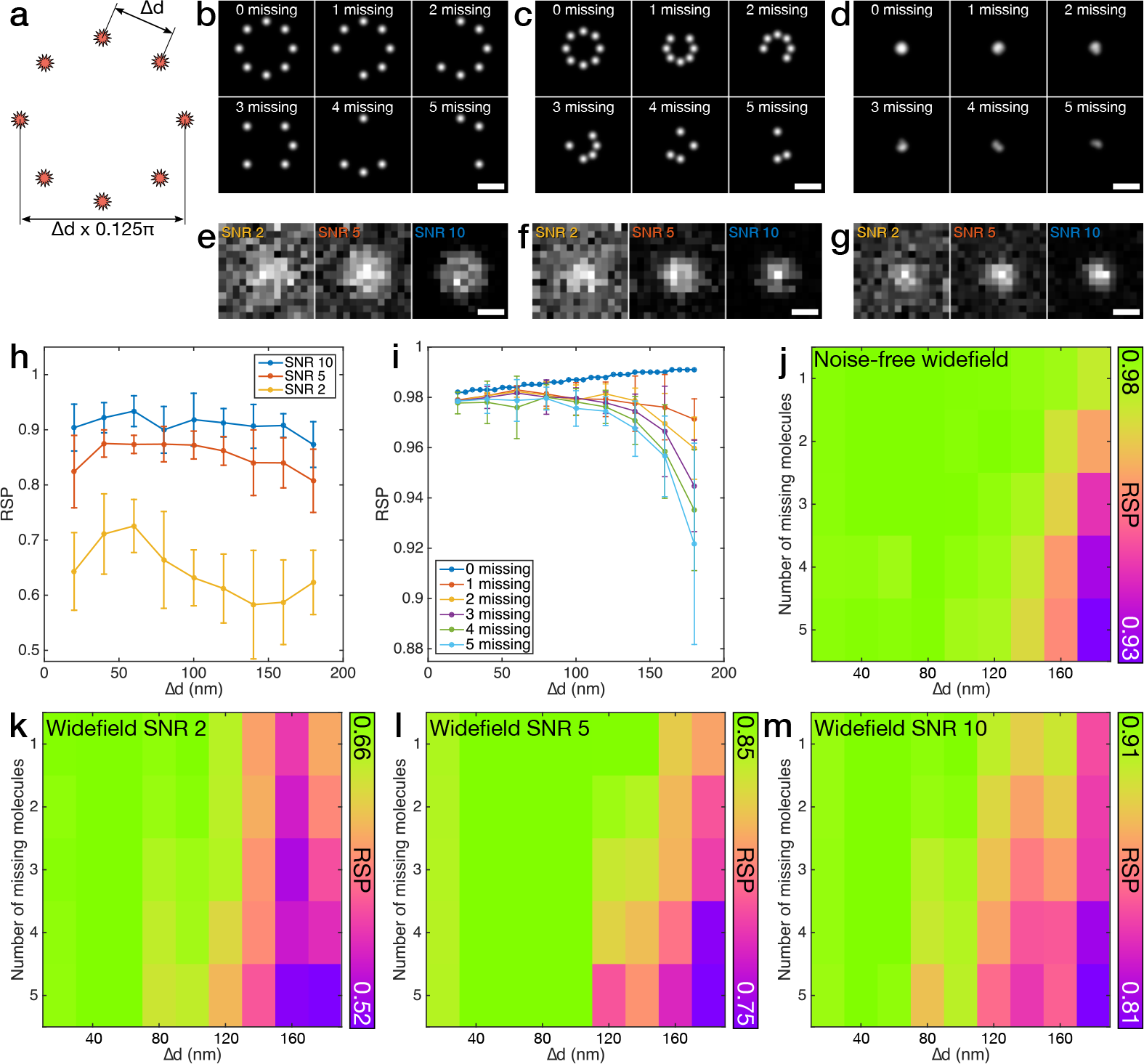
Resolution and signal-to-noise sensitivity of SQUIRREL. a) Geometry of an 8-molecule ring structure for simulations, where Δd indicates the separation between adjacent molecules. b-d) Representative simulated super-resolution images with varying numbers of randomly removed molecules from the ring structure with Δd values of b) 160 nm, c) 100 nm, d) 20 nm, rendered with Gaussian distributions of *σ=20* nm. Scale bars = 200 nm. e-g) Representative simulated widefield images of rings with no molecules missing at three different signal-to-noise ratios (SNR) for Δd values of e) 160 nm, f) 100 nm, g) 20 nm, where the widefield PSF is approximated to a Gaussian distribution of σ=135 nm. Scale bars = 200 nm. h) RSP values generated by SQUIRREL for simulated super-resolution images with no molecules missing with different SNR widefield images used as the reference. Error bars represent standard deviation of n=10 repeats. i) RSP values generated by SQUIRREL for super-resolution images with varying numbers of molecules missing when a noise-free widefield image is used as the reference. Error bars represent standard deviation of n=100 repeats. j-m) Mean RSP values mapped for number of molecules removed against Δd for widefield reference images with j) no noise (n=100), k) SNR = 2 (n=10), l) SNR = 5 (n=10), m) SNR = 10 (n=10).

‘Perfect’ super-resolution images were simulated for different sized rings with all 8 molecules present, and artefactual superresolution images were generated through random removal of various numbers of molecules from the ring (Fig. S4b-d). Widefield images with all 8 molecules present were also generated to serve as SQUIRREL reference images. These either feature no noise, or have one of three different signal-to-noise ratios (SNR) (Fig. S4e-g). When the super-resolution image contained no artefacts, the RSP values were dictated by the SNR of the reference image, with higher SNR yielding higher RSP values (Fig. S4h). The RSP does not significantly vary when changing the size of the ring structure for a constant SNR. Next, the sensitivity of SQUIRREL was assessed for artefactual super-resolution images with up to 5 molecules absent when compared with a noise-free reference image with all 8 molecules present (Fig. S4i). When there were no molecules removed from the super-resolution image, the RSP steadily increased with Δd; however, when molecules were removed from the superresolution image the opposite relationship was observed, with the RSP decreasing at larger values of Δd. This relationshipbecame more pronounced with greater numbers of removed molecules (i.e. more artefactual images). For Δd less than 100 nm, the RSP values were not significantly different to the perfect super-resolution images regardless of the number of molecules absent from the structure. This is due to the lack of resolution support in the widefield image. However, the RSP indicated the presence of artefacts at all larger Δd values. The larger error bars associated with the RSP values for increasingly artefactual super-resolution images are due to the random nature of multiple molecule removals. The relationship between the degree of image artefacts (i.e. number of missing molecules) and the size of the structure (i.e. Δd) is displayed in Fig. S4j. The combined effect of reference image SNR, extent of super-resolution image defects and structure size is plotted similarly in Fig. S4k-m. As in Fig. S4h RSP values are consistently lower for low SNR, and as in Fig. S4j there is higher variation in RSP values for more extreme defects and larger structures. Based on these results, we conclude that SQUIRREL is sufficiently sensitive to describe defects occurring on a scale down to ∼150 nm, with even smaller scales possible for high SNR.

**Fig. S5.**
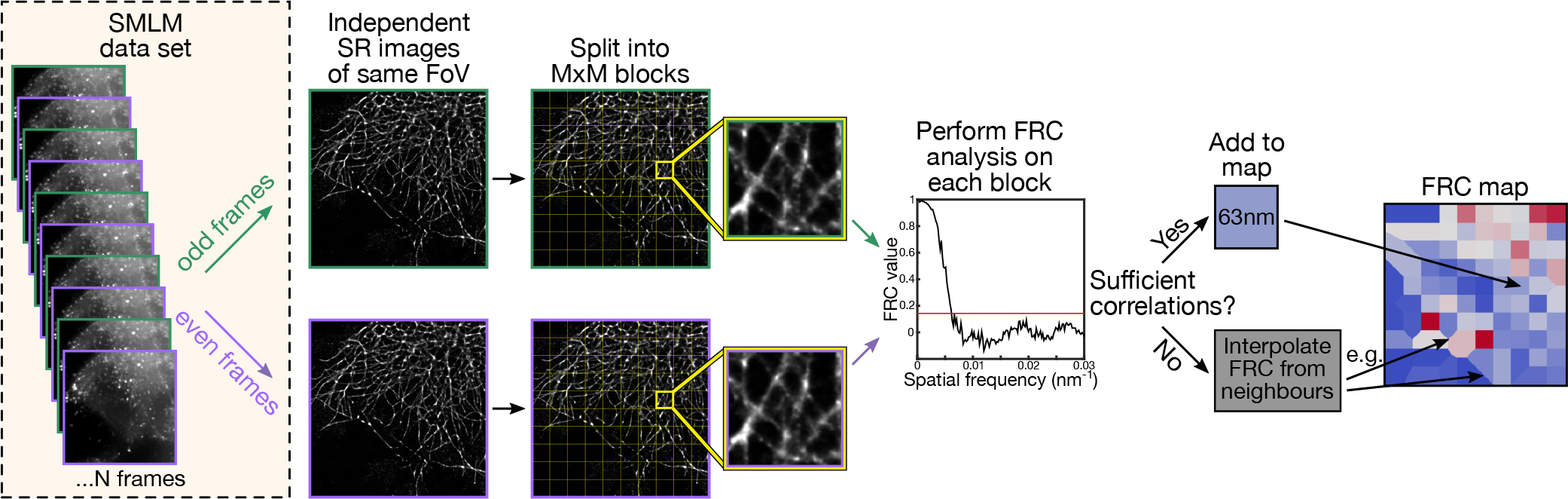
FRC mapping. Resolution mapping by FRC is carried out by inputting two independent super-resolution images of the same field-of-view imaged under the same conditions. For the case of SMLM data, these two such images can be generated by separating the data in half, for example by having odd and even frames producing two independent super-resolution reconstructions. Within the algorithm, the two data reconstructions are divided into blocks and for each block the FRC value is calculated as described in (14). If sufficient correlations for FRC resolution estimation is found in a block, an equivalent region is colour coded to this value in an FRC resolution map. If not, the equivalent region is colour coded according to natural neighbour interpolation (15).

## Supplementary Note 5: Multi-Image Elastic Registration

In multi-image elastic registration, the image is divided into a user-selected number of blocks (similarly to Fig. S5). For each block, an optimal Δ*x*, Δ*y* translation is calculated such that local similarity between the block in the image currently being registered and the block in the reference image is maximised. This calculation is done by cross-correlation, in the same manner described in Sup. Note 2. If a block does not support sufficient correlation against its given reference block, *Δx, Δy* will be predicted by interpolation based on the values of neighbouring blocks. An elastic translation matrix is then calculated by inverse distance weighting (16) and applied to the corresponding image by bi-cubic spline interpolation (8).

**Fig. S6.**
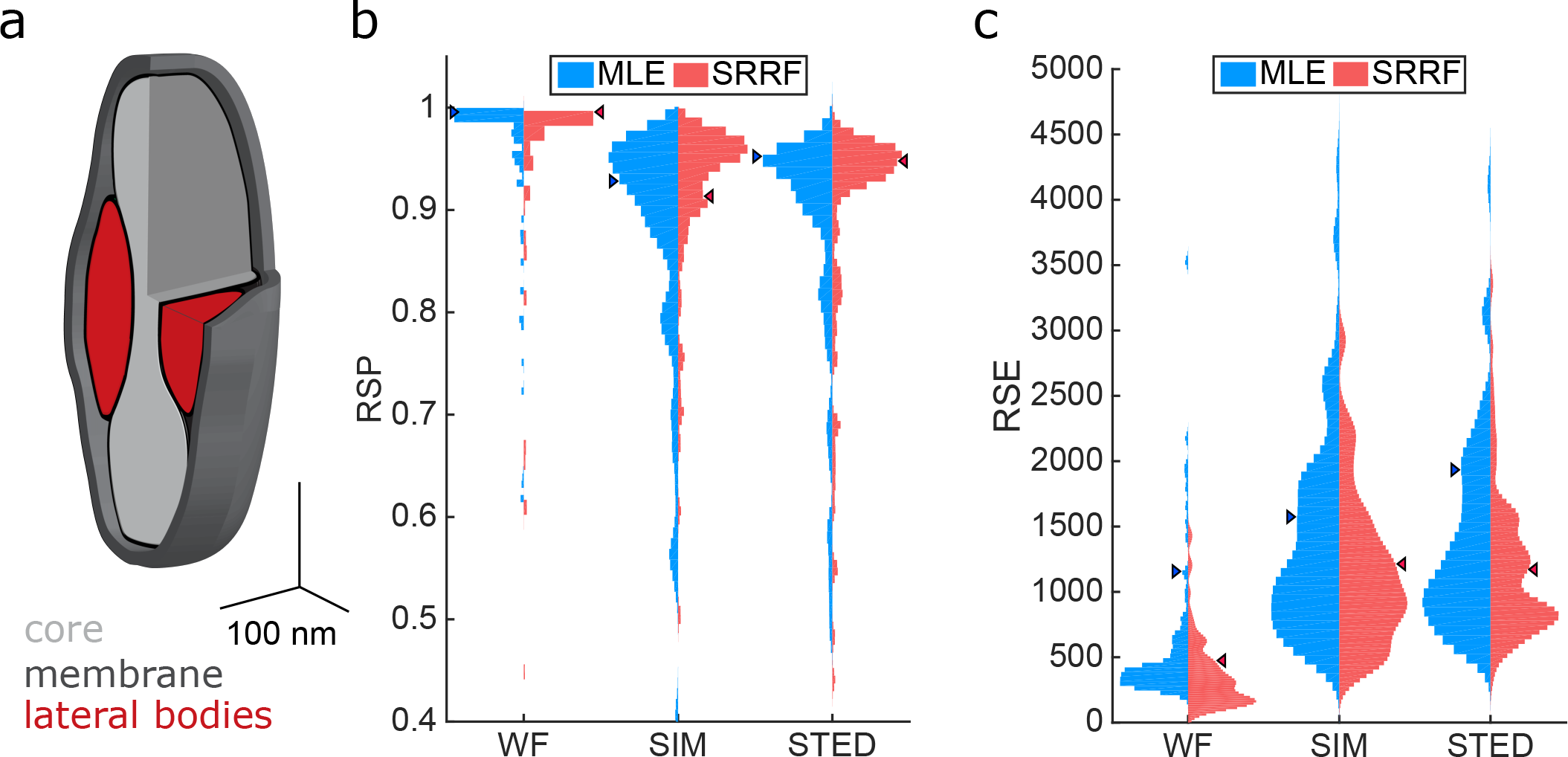
Comparative image quality between super-resolution methods while imaging vaccinia virus lateral bodies. a) Schematic of vaccinia structure, red highlights the lateral bodies with the remaining major structural components shown in grey, b) Violin plots of RSP values when comparing MLE and SRRF reconstructions against widefield, SIM and STED reference images for 90 individual viruses from the images in Fig, 3a, Arrowheads indicate values for the example virus in Fig, 3b, c) As in **b**, but RSE values

## Supplementary Note 6: Image Fusion

Image fusion provides a super-resolution estimate merging the best traits of multiple super-resolution reconstructions through a weighted average. For this purpose, the fusion process weights the local information of each reconstruction according to its local error to achieve a final estimate of higher quality than any of its components. The process starts by the calculation of *R(n, x, y)* as the root-mean-square-error (RMSE) of each reconstruction *n* for a small 3x3 window (Eq. S7). Here *M(n, x, y)* corresponds to the error map matrix (Eq. 4) for the n-th super-resolution reconstruction provided. Next a matrix R_max_(x, *y)* is also calculated, containing the maximum value of *R(n, x, y)* for each pixel across *N* total reconstructions (Eq. S8). A weight matrix for each super-resolution reconstruction can then be calculated through Eq. S9 (note the truncation of the denominator to 1 in order to avoid a zero-division). For the *W(n, *x*, *y*)* calculation we have chosen to use the RMSE over the 3x3 as described in Eq. S7 instead of *M(n, x, y)* directly to provide local error smoothing, minimising sharp pixel-wise changes between individual dominant pixels from each reconstruction. The final fusion image *I*_SF_ *(x, y)* is generated through a weighted average (Eq. S10).

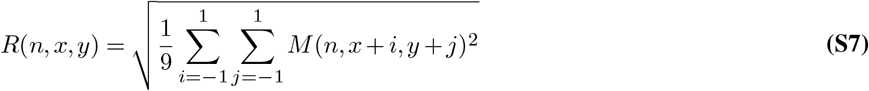

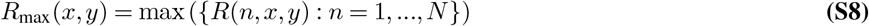

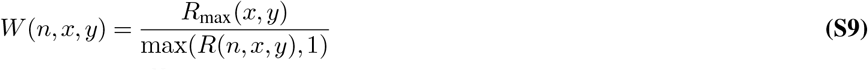

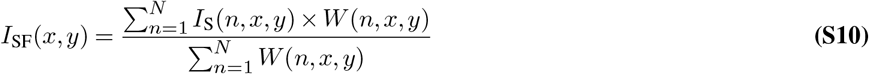

**Fig. S7.**
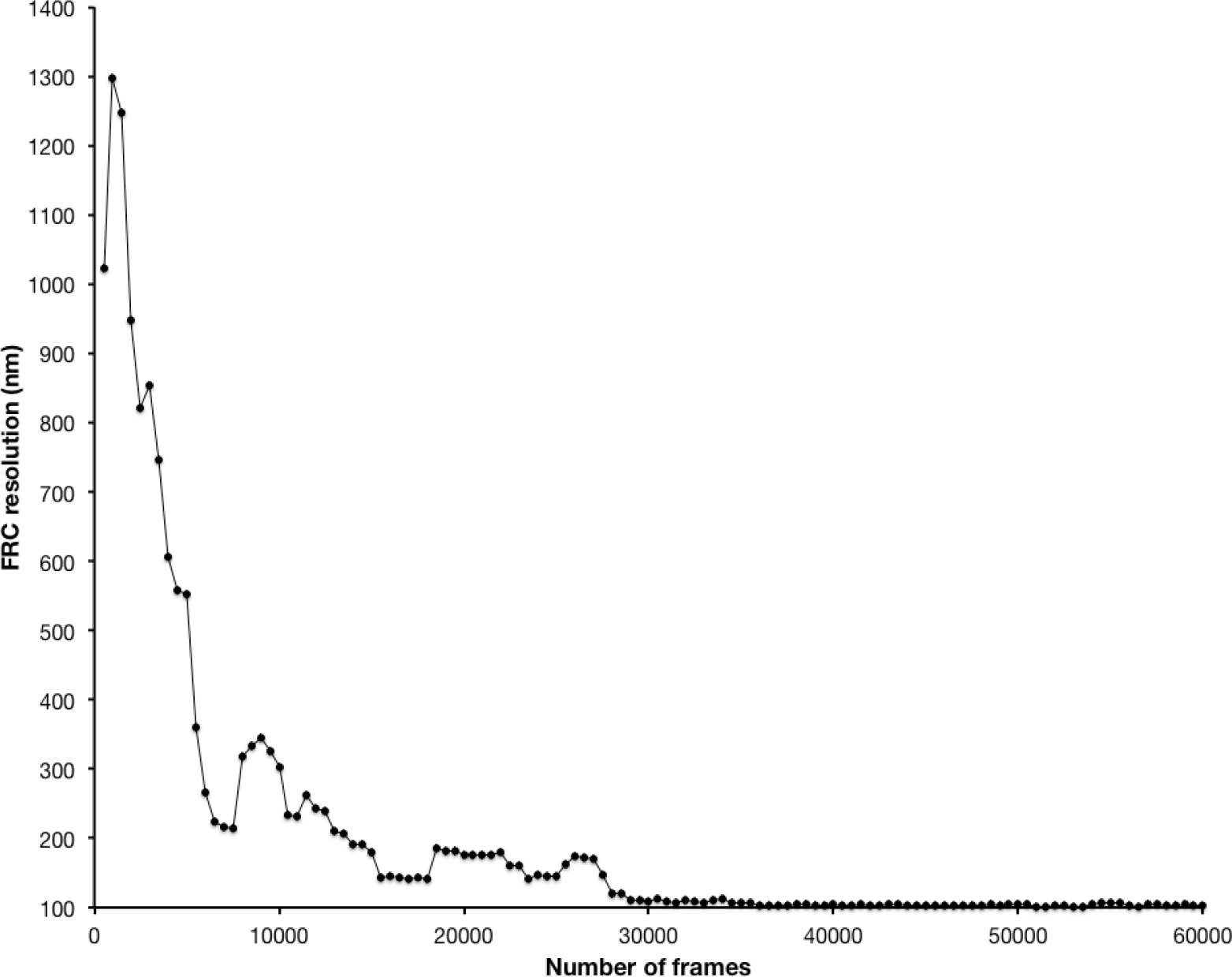
Fourier Ring Correlation resolution for neuronal actin rings. Average FRC values for reconstructions obtained with an accumulative number of frames, in the same manner as Fig. 5e-f. ⋅

## Supplementary Methods

**SIM actin imaging.** For comparison of different SNR ratios, FluoCells prepared slide #2 (Invitrogen) with BPAE cells stained with Texas Red-X phalloidin was imaged on a Zeiss Elyra PS.1 system, using a 63 × NA 1.4 objective with additional 1.6× magnification for SIM and widefield acquisition. Low SNR images were acquired with a 561 nm laser at 0.05 % laser power, using 100 ms exposure time, and 5 grid rotations. High SNR images were acquired with a 561 nm laser at 5 % laser power, 100 ms exposure, 5 grid rotations. Widefield images were acquired with a 561 nm laser at 0.2 % laser power, 100 ms exposure time. SIM reconstructions were generated with the Zeiss Elyra Zen software using automatic settings.

**Simulation of structures with out-of-focus information.** For assessing the impact of out-of-focus fluorescence on SQUIRREL analysis, a test structure was simulated consisting of three semicircles of radius 500 nm and axial tilt ranging from –750 nm to +750 nm. A widefield representation of this structures was produced via convolution with a 3D PSF generated using the ImageJ PSF Generator plugin (6) with the following settings: Born and Wolf optical model, numerical aperture 1.4, wavelength 640 nm, z-range 1500 nm, z-step size 2 nm. Single molecule blinking data sets were generated with custom-written simulation software with the same PSF used for rendering molecule appearances, and were binned into 100nm ‘camera’ pixels. This was performed for both the defect-free structure and an artefactual equivalent where 100 nm stretches of the structure had been deleted. These data sets were analysed using weighted-least-squares fitting in ThunderSTORM (13) and rendered with 20 nm Gaussian distributions.

